# Single-molecule sequencing and conformational capture enable *de novo* mammalian reference genomes

**DOI:** 10.1101/064352

**Authors:** Derek M. Bickhart, Benjamin D. Rosen, Sergey Koren, Brian L. Sayre, Alex R. Hastie, Saki Chan, Joyce Lee, Ernest T. Lam, Ivan Liachko, Shawn T. Sullivan, Joshua N. Burton, Heather J. Huson, Christy M. Kelley, Jana L. Hutchison, Yang Zhou, Jiajie Sun, Alessandra Crisà, F. Abel Ponce De León, John C. Schwartz, John A. Hammond, Geoffrey C. Waldbieser, Steven G. Schroeder, George E. Liu, Maitreya J. Dunham, Jay Shendure, Tad S. Sonstegard, Adam M. Phillippy, Curtis P. Van Tassell, Timothy P.L. Smith

**Affiliations:** Animal Genomics and Improvement Laboratory, ARS USDA, Beltsville, Maryland, USA 20705; National Human Genome Research Institute, Bethesda, USA 21702; Department of Biology, Virginia State University, Petersburg, Virginia, USA 23806; BioNano Genomics, San Diego, California, USA 92121; Department of Genome Sciences, University of Washington School of Medicine, Seattle, WA 98195; Phase Genomics, 4000 Mason Road, Suite 225, Seattle, WA 98195; Department of Animal Science, Cornell University, Ithaca, New York, USA 14853; Genetics, Breeding and Animal Health Research, ARS USDA, Clay Center, Nebraska, USA 68933; South China Agricultural University, 483 Wushan Rd, Tianhe, Guangzhou, Guangdong, China; CRA Agricultural Research Council, Research Centre for Meat Production and Genetic Improvement, Rome, Italy; Department of Animal Science, University of Minnesota, St. Paul, MN, USA 55108; Livestock Viral Disease Programme, The Pirbright Institute, Woking, GU24 0NF, UK; Warmwater Aquaculture Research Unit, ARS USDA, Stoneville, Mississippi, USA 38776; Howard Hughes Medical Institute, Seattle WA 98195; Recombinetics, Inc. 1246 University Ave W #301, St. Paul, MN 55104

## Abstract

The decrease in sequencing cost and increased sophistication of assembly algorithms for short-read platforms has resulted in a sharp increase in the number of species with genome assemblies. However, these assemblies are highly fragmented, with many gaps, ambiguities, and errors, impeding downstream applications. We demonstrate current state of the art for *de novo* assembly using the domestic goat (*Capra hircus*), based on long reads for contig formation, short reads for consensus validation, and scaffolding by optical and chromatin interaction mapping. These combined technologies produced the most contiguous *de novo* mammalian assembly to date, with chromosome-length scaffolds and only 663 gaps. Our assembly represents a >250-fold improvement in contiguity compared to the previously published *C. hircus* assembly, and better resolves repetitive structures longer than 1 kb, supporting the most complete repeat family and immune gene complex representation ever produced for a ruminant species.

## Introduction

A finished, accurate reference genome provides an essential component for advanced genomic selection of productive traits in agriculturally relevant plant and animal species^1–3^. Thus, better genome finishing technologies will be of immediate benefit to researchers of these organisms. Substantial progress has been made in methods for generating contigs from whole genome shotgun (WGS) sequencing; yet finishing genomes remains a labor-intensive process that is unfeasible for most large, highly repetitive genomes. For example, the successful production of the human reference genome assembly draft in 2001^4^ was followed by three years of intensive curation by 18 individual institutions^5^, including BAC primer walking experiments to close gaps and manual assembly inspection. This effort was rewarded with the best reference genome assembly for a mammalian species, which presently (version GRCh38) contains only 832 gaps whose content primarily corresponds to heterochromatin. The advent of massively parallel DNA sequencing technologies in 2005 democratized sequencing, allowing hundreds of reference genomes to be generated. However, equivalently high-throughput methods for finishing were not available, and these assemblies remain highly fragmented^6^.

Repeats pose the largest challenge for reference genome assembly, and much effort has been devoted to resolving the ambiguous assembly gaps caused by repetitive sequence^7^. The process of ordering and orienting assembled contigs around such gaps is known as scaffolding,^8, 9^, mate-pair scaffolding^9, 10^, long-read scaffolding^11^, compartmentalized shearing and barcoding^12^, chromatic interaction mapping (Hi-C)^13^, and optical mapping^14^. Of these methods, both Hi-C and optical mapping provide relatively inexpensive and high-resolution mapping data that can be useful for scaffolding^15–19^. Hi-C is an adaptation of the chromosome conformation capture (3C) methodology^20^ that can identify long-range chromosome interactions in an unbiased fashion, without *a priori* target site selection. By observing long-range consensus interactions, whose frequency decays rapidly based on linear distance on the same chromosome, Hi-C data can be used to scaffold assembled contigs to the scale of full chromosomes^15^. In comparison, optical mapping technologies observe the linear separation of small DNA motifs (often restriction enzyme recognition sites^19^ or nickase sites^21^), which can provide sufficient contextual information to scaffold assembled contigs^22^ or correct existing reference assemblies^23^. Both optical mapping^21^ and Hi-C^15^ yield excellent scaffold continuity metrics ^15, 17, 18, 24^. However, both methods are limited in their ability to map short contigs and so Hi-C or optical map scaffolds based on fragmented short-read assemblies often remain incomplete^25^.

Single-molecule sequencing^26^ is now capable of producing reads tens of kilobases in size, albeit with relatively high error rate. For example, the PacBio RSII sequencing platform can achieve an average read length of 14 kb or larger and maximum read lengths exceeding 60 kb^27^.This platform is now routinely used to reconstruct complete bacterial genomes^28, 29^ and has been recently applied to eukaryotic genomes ^27, 30, 31^. Because the maximum read length exceeds the size of most repetitive structures, long-read sequencing is theoretically capable of assembling near-complete mammalian chromosomes, but this would require sequencing to an impractical depth to collect enough of these very long reads to span all of the large repeats. As a result, published long-read mammalian assemblies still comprise thousands of pieces ^27, 30^. Until the long read platforms can regularly produce average lengths of >30 kb, or become cheap and efficient enough to consider deep coverage, combinations of long-read assemblies and long-range scaffolding techniques represent the most efficient approach to produce a finished—or near finished—reference assembly. Indeed, a proof-of-concept study using PacBio single-molecule real-time (SMRT) sequencing and BioNano Genomics optical mapping recently assembled a human genome *de novo* into 4,007 contigs and 202 scaffolds that covered the entire reference assembly^31^.

Here we present a near-finished reference genome for the domestic goat (*Capra hircus*) generated via an improved assembly strategy using a combination of long-read single-molecule sequencing, high-fidelity short read sequencing, optical mapping, and Hi-C-based chromatin interaction maps. The goat is one of four major ruminant species used for human food production and is an ideal species for comparative and population genomic studies. Because the developing world contains approximately 95% of the world goat population (http://www.fao.org/docrep/009/ah221eZAH221E13.htm), improvement of goat meat and milk production efficiency in these regions is one of the critical components to meeting global food security challenges due to climate change and expanding population sizes. There is supporting evidence that all breeds of goats are derived from a single wild ancestor, the bezoar^32^, unlike cattle, which are derived from two different sub-species^33^. Due to this singular domestication event, creation of a polished reference genome for goat could enable easier identification of adaptive variants in sequence data from descendent breeds. However, the most recent goat
assembly was generated via only short-read sequencing and optical mapping, and remains heavily fragmented as a result^18^. Our new assembly strategy is cost effective compared to past finishing approaches, achieves unsurpassed continuity and accuracy, and provides a new standard reference for ruminant genetics.

## Results

### *de novo* assembly of a *Capra hircus* reference genome

We selected an individual goat of the San Clemente breed that displayed a high degree of homozygosity for sequencing in order to minimize heterozygous alleles and simplify assembly. Data was collected from this single individual using a combination of four technologies: singlemolecule real-time sequencing (PacBio RSII), paired-end sequencing (Illumina HiSeq), optical mapping (BioNano Genomics Irys), and Hi-C proximity guided assembly (Phase Genomics). Assembly of these complementary data types proceeded in a stepwise fashion (Methods), producing progressively improved assemblies (Table 1). Initial contigs were first assembled from the PacBio data alone, resulting in a contig NG50 of 4.16 Mbp (NG50: contig size such that half of the haploid genome is accounted for by contigs of this size or greater). These PacBio contigs were then further corrected, joined, ordered, and oriented using the Irys optical mapping data, increasing contig size and forming an initial set of scaffolds, which were then combined with Hi-C data to further extend scaffolds and cluster the chromosomes. Scaffold gaps were then filled where possible using mapped single-molecule data, and the final assembly was polished to achieve high consensus accuracy using mapped Illumina data. The resulting reference assembly, named ARS1, totals 2.92 Gbp of sequence with a contig NG50 of 19 Mbp and a scaffold NG50 of 87 Mbp, and was further validated via statistical methods and comparison to a radiation hybrid (RH) map^34^ and previous assemblies (Supplementary Note 1).

### Scaffolding method comparisons

We compared *de novo* optical map and Hi-C scaffolds to our validated reference assembly in order to evaluate the independent performance of the two scaffolding methods. Optical map scaffolding of PacBio contigs alone resulted in an assembly of 842 scaffolds containing 91.5% of the total assembly length with a scaffold NG50 of 13.4 Mbp and 34 confirmed scaffold conflicts. Optical map (Supplementary Fig. S1) scaffold sizes were primarily limited by double-strand breaks caused by close proximity of *Nt.Bsql* sites on opposing DNA strands, as reported previously^21^. However, the differences in the error-profile between the PacBio contigs and optical map caused subsequent scaffolding to synergistically increase final scaffold NG50 values three-fold (Table 1). This is likely due to the PacBio’s ability to sequence through shorter, low-complexity repeats, combined with the optical map’s ability to span larger segmental duplications. In comparison, scaffolding of PacBio contigs with Hi-C data yielded 31 scaffolds containing 89.6% of the total assembly length (Supplementary Fig. S2 and Supplementary Table S1). These scaffolds had an NG50 four times larger than the scaffolds generated by optical mapping, but had a relatively higher rate of mis-oriented contigs when compared to the RH map^34^ (i.e. PacBio contigs assigned to the wrong DNA strand within a scaffold). Analysis of the mis-oriented contigs revealed that orientation error was proportional to the density of Hi-C restriction sites in the contig (Supplementary Table S2), and so future assembly projects should choose restriction enzymes with shorter recognition sites (or DNase Hi-C^35^) to improve Hi-C link density and the associated orientation error rate.

**Table 1.**
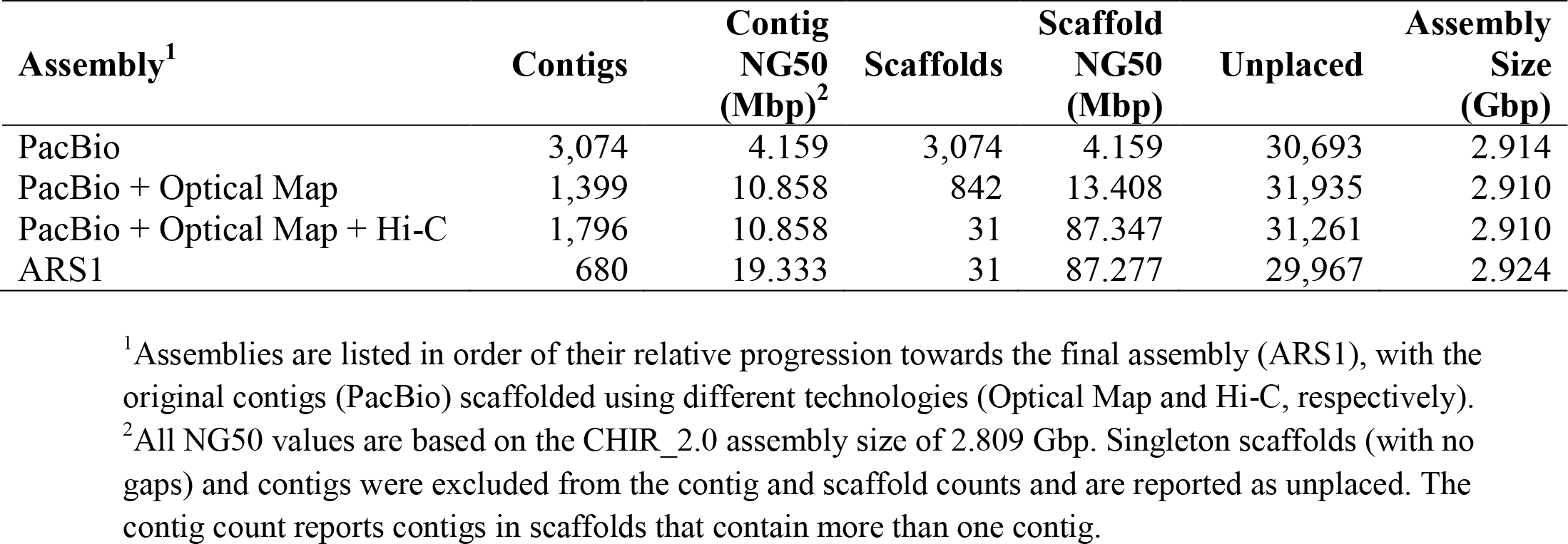
Assembly statistics

| Assembly<sup>1</sup> | Contigs | Contig<br>NG50<br>(Mbp)<sup>2</sup> | Scaffolds | Scaffold<br>NG50<br>(Mbp) | Unplaced | Assembly<br>Size<br>(Gbp) |
| --- | --- | --- | --- | --- | --- | --- |
| PacBio | 3,074 | 4.159 | 3,074 | 4.159 | 30,693 | 2.914 |
| PacBio + Optical Map | 1,399 | 10.858 | 842 | 13.408 | 31,935 | 2.910 |
| PacBio + Optical Map + Hi-C | 1,796 | 10.858 | 31 | 87.347 | 31,261 | 2.910 |
| ARS1 | 680 | 19.333 | 31 | 87.277 | 29,967 | 2.924 |

<sup>1</sup>Assemblies are listed in order of their relative progression towards the final assembly (ARS1), with the original contigs (PacBio) scaffolded using different technologies (Optical Map and Hi-C, respectively).
<sup>2</sup>All NG50 values are based on the CHIR\_2.0 assembly size of 2.809 Gbp. Singleton scaffolds (with no gaps) and contigs were excluded from the contig and scaffold counts and are reported as unplaced. The contig count reports contigs in scaffolds that contain more than one contig.

Ultimately, we found that sequential scaffolding with optical mapping data followed by Hi-C data yielded an assembly with the highest continuity and best agreement with the RH map (Fig. 1). This provided the Hi-C step with initial scaffolds, which could be more confidently arranged into chromosomes owing to the greater number of Hi-C links between scaffolds than between the relatively shorter contigs. Post-scaffolding, we then applied a series of error-correction methods to fill gaps and remove artifacts from our assembly^36, 37^. This resulted in a final assembly with 31 chromosome-scale scaffolds, 652 unplaced scaffolds, 29,315 unplaced contigs, 663 gaps, and an estimated QV of 34.5 (Fig. 2, Supplementary Note 1)^38^. Excluding small or repetitive contigs that could not be placed on the RH map, we measured 99.8% agreement between our ARS1 assembly and the RH map (1529/1533 correctly scaffolded contigs), leaving just four regions of major disagreement that will require further investigation (Fig. 3). Considering that ARS1 contains 31 scaffolds with only 663 gaps covering 30 of the 31 haploid, acrocentric goat chromosomes^39^ (excluding only the Y chromosome), our assembly statistics compare favorably with the current human reference (GRCh38) with 24 scaffolds, 169 unplaced/unlocalized scaffolds, and 832 gaps in the primary assembly (http://www.ncbi.nlm.nih.gov/projects/genome/assembly/grc/).

**Figure 1:**
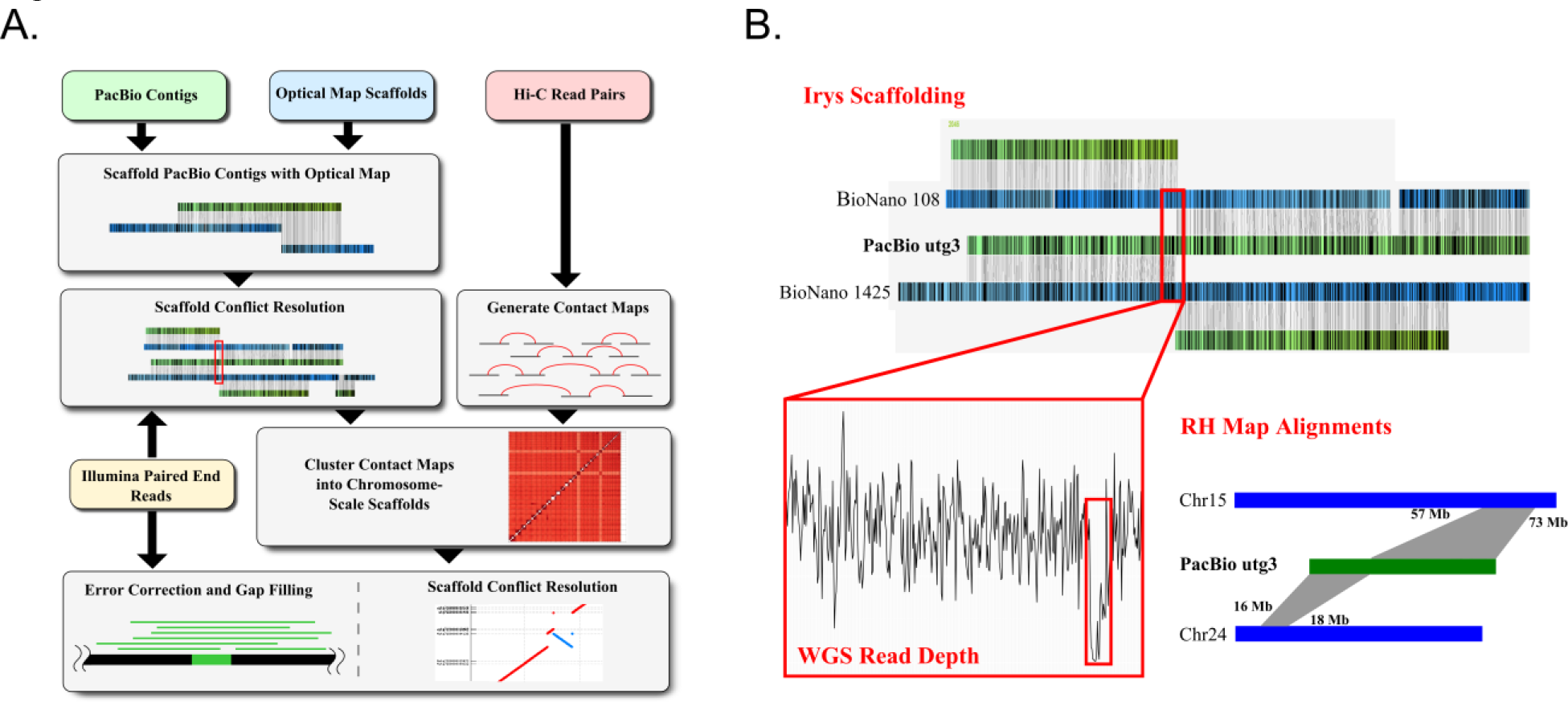
Assembly schema for producing chromosome-length scaffolds. (A) Four different sets of sequencing data (long-read WGS, Hi-C data, optical mapping and short-read WGS) were produced in order to generate the goat reference genome. A tiered scaffolding approach using optical mapping data followed by Hi-C proximity guided assembly produced the highest quality genome assembly. (B) In order to correct mis-assemblies resulting from contig-or scaffold-errors, a consensus approach was used. An example from the initial optical mapping dataset is shown in the figure. A scaffold fork was identified on contig 3 (a 91 Mbp length contig) from the optical mapping data. Subsequent short-read WGS data and an existing RH map for goat showed signatures that there was a mis-assembly near the 13^th^ megabase of the contig, so it was split at this region.

**Figure 2:**
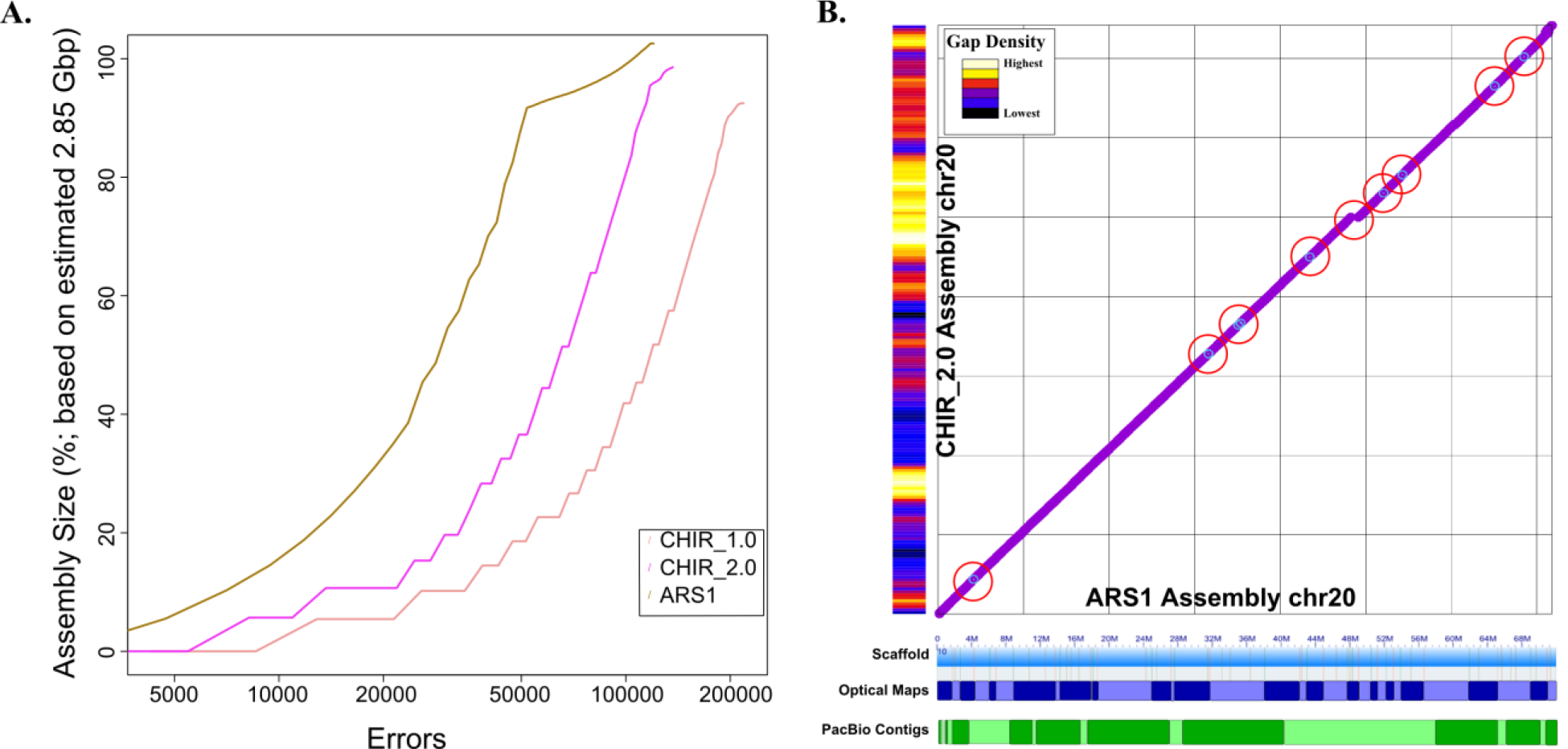
Assembly benchmarking comparisons reveal high degree of assembly completion. (A) Feature response curves (FRC) showing the error rate as a function of the number of bases in each assembly (CHIR_1.0, CHIR_2.0 and ARS1) and each scaffold test (Hi-C and BioNano). (B) Comparison plots of chromosome 20 sequence between the ARS1 and CHIR_2.0 assemblies reveal several small inversions (light blue circles) and a small insertion of sequence (break in continuity) in the ARS1 assembly. Red circles highlight 9 of the aforementioned inversions and the insertion of sequence in our assembly. The ARS1 assembly contains only 10 gaps on this chromosome scaffold whereas CHIR_2.0 has 5,651 gaps on the same chromosome assembly (gap density histogram on the Y axis). ARS1 optical map scaffolds and Pacbio contigs represented on the X axis as alternating patterns of blue and green shades, respectively, showing the tiling path that comprises the entire single chromosome scaffold.

**Figure 3:**
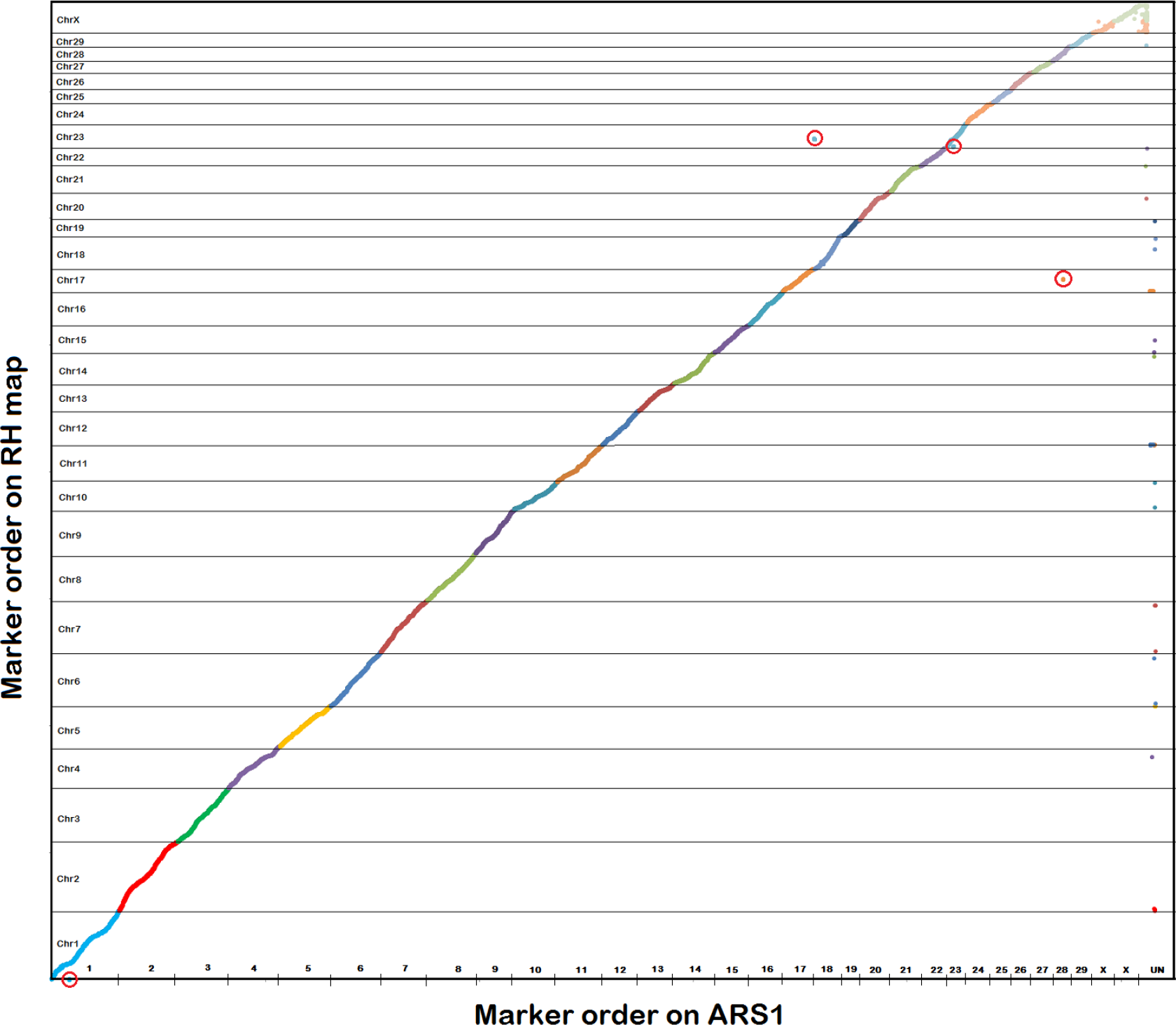
RH probe map shows excellent assembly continuity. ARS1 RH probe mapping locations were plotted against the RH map order. Each ARS1 scaffold corresponds to an RH map chromosome with the exception of X which is composed of two scaffolds. Red circles highlight two intrachromosomal (on chrs 1 and 23) and two interchromosomal mis-assemblies (on chrs 18 and 17) in ARS1 that were difficult to resolve.

### Assembly benchmarking and comparison to reference

We used several methods to quantify the improvement of our assembly over previous reference assemblies built only upon short-read sequencing and scaffolding techniques. Short paired-end read sequences from the Black Yunan goat CHIR_1.0 reference assembly^18^ were aligned to CHIR_1.0, CHIR_2.0 and our ARS1 assembly for a reference-free measure structural correctness^40, 41^ (Supplementary Note 2). These alignments revealed that CHIR_2.0 was a general improvement over CHIR_1.0, resulting in fewer putative deletions (2,735 vs. 10,256) and duplications (115 vs. 290) compared to the previous reference assembly; however, CHIR_2.0 also contains 50-fold more putative inversions than CHIR_1.0 (215 vs. 4; Supplementary Table S3). Our ARS1 assembly was found to be a further improvement over CHIR_2.0, with fourfold fewer deletions and 50-fold fewer inversions identified. This is particularly notable given that the Black Yunan data was not used in our assembly and yet our assembly is more consistent with this data than the CHIR_1.0 and CHIR_2.0 assemblies themselves. Thus, this independent validation indicates that ARS1 corrects numerous errors present in CHIR_2.0 (Fig. 2).

We also assessed the quantity and size of gaps in each respective assembly (Supplementary Table S5). While the CHIR_2.0 reference was able to fill 62.4% of CHIR_1.0 gap sequences (160,299 gaps filled), our assembly filled 94.6% of all CHIR_1.0 gaps (242,888 gaps filled). The remaining CHIR_1.0 gaps (13,853) had flanking sequence that mapped to two separate chromosomes in our assembly, indicating potential false gaps due to errors in the CHIR_1.0 assembly. Sequence alignments from our San Clemente reference animal as well as the RH map and CHIR_2.0 agreed with our assembly in these locations (Methods), confirming the CHIR_1.0 errors. In total, our assembly contains only 663 sequence gaps (larger than 3 bp) in the chromosomal and unassigned scaffolds split among gaps of known (48 inferred from optical mapping distances) and unknown (615 Hi-C scaffold joining) sizes. Finally, compared to CHIR_2.0, ARS1 has 1000-fold fewer ambiguous bases and a two point higher BUSCO score^42^ (82% vs 80%, respectively).

### Improvement in genetic marker tools and functional annotation

We quantified the benefit of our approach over short-read assembly methods with respect to genome annotation and downstream functional analysis. Chromosome-scale continuity of the ARS1 assembly was found to have appreciable positive impact on genetic marker order for the existing *Capra hircus* 52k SNP chip^3^ (Supplementary Table S4). Of the 1,723 SNP probes currently mapped to the unplaced contigs of the CHIR_2.0 assembly, we identified chromosome locations for 1,552 (90.0%) of the markers and identified 26 additional, low call-rate SNP markers as having ambiguous mapping locations in our assembly^3^. This finding suggests these markers were unknowingly targeting repeat sequences and provides an explanation as to why they were poor performers on the chip. In order to identify improvements in gene annotation resulting from our method of assembly, we focused on gap regions from previous assemblies that intersected with newly annotated gene models (Methods). We found that 3,495 of our gene models had at least one CHIR_2.0 gap within either an exon or an intron that was filled by our assembly (Supplementary Table S5). We also identified 1,926 predicted exons that contained gaps in CHIR_1.0 and CHIR_2.0 but were resolved by our assembly (Fig. 4a). Annotation of repetitive immune gene regions revealed that complete complements of the leukocyte receptor complex (LRC) and natural killer cell complex (NKC) were contained within single autosome scaffolds in our assembly (Fig. 5). These regions are particularly difficult to assemble with short-read technologies because they are highly polymorphic and repetitive^43^, with gene content being largely species specific, so it is not surprising that these regions are fragmented among separate scaffolds in the CHIR_1.0 and CHIR_2.0 assemblies (Supplementary Note 3 and Supplementary Figs. S3, S4, and S5).

**Figure 4:**
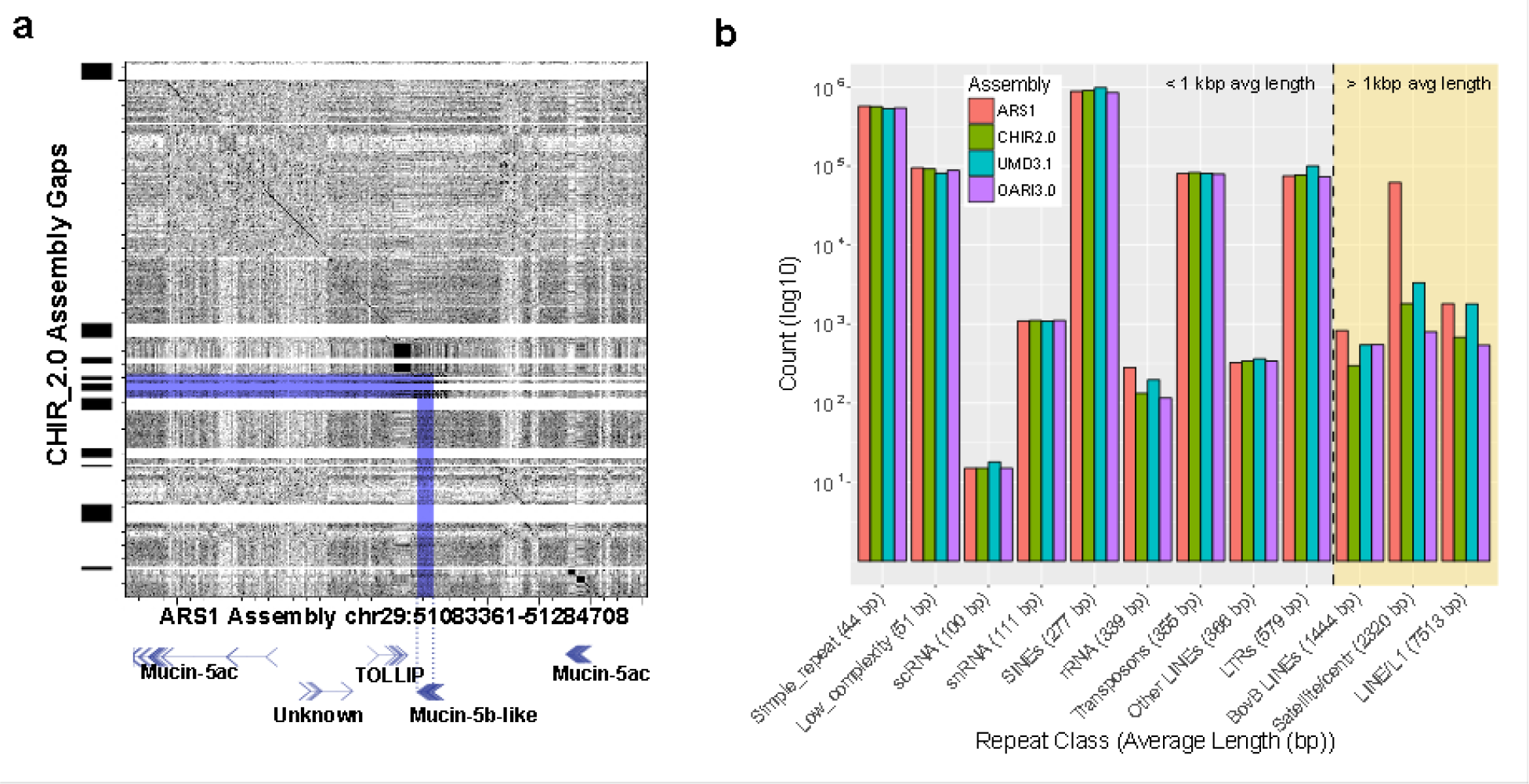
Long-read assembly with complementary scaffolding resolves gap regions (A) and long repeats (B) that cause problems for short-read reference annotation. (A) A region of the Mucin gene cluster was resolved by long-read assembly, resulting in a complete gene model for Mucin-5b-Like that was impossible due to two assembly gaps in the CHIR_2.0 assembly. (B) Counts of repetitive elements that had greater than 75% sequence length and greater than 60% identity with RepBase database entries for ruminant lineages. With the exception of the rRNA cluster (which is present in many repeated copies in the genome), the CHIR_2.0 reference contained a full complement of shorter repeat segments that were also present in our assembly. However, repeats that were larger than 1 kb (highlighted by the yellow background) were present in higher numbers in our assembly due to our ability to traverse the entire repetitive element’s length. Our assembly resolves repeats better than the multi-million dollar cattle assembly^44^ across multiple repeat classes >1kbp.

**Figure 5:**
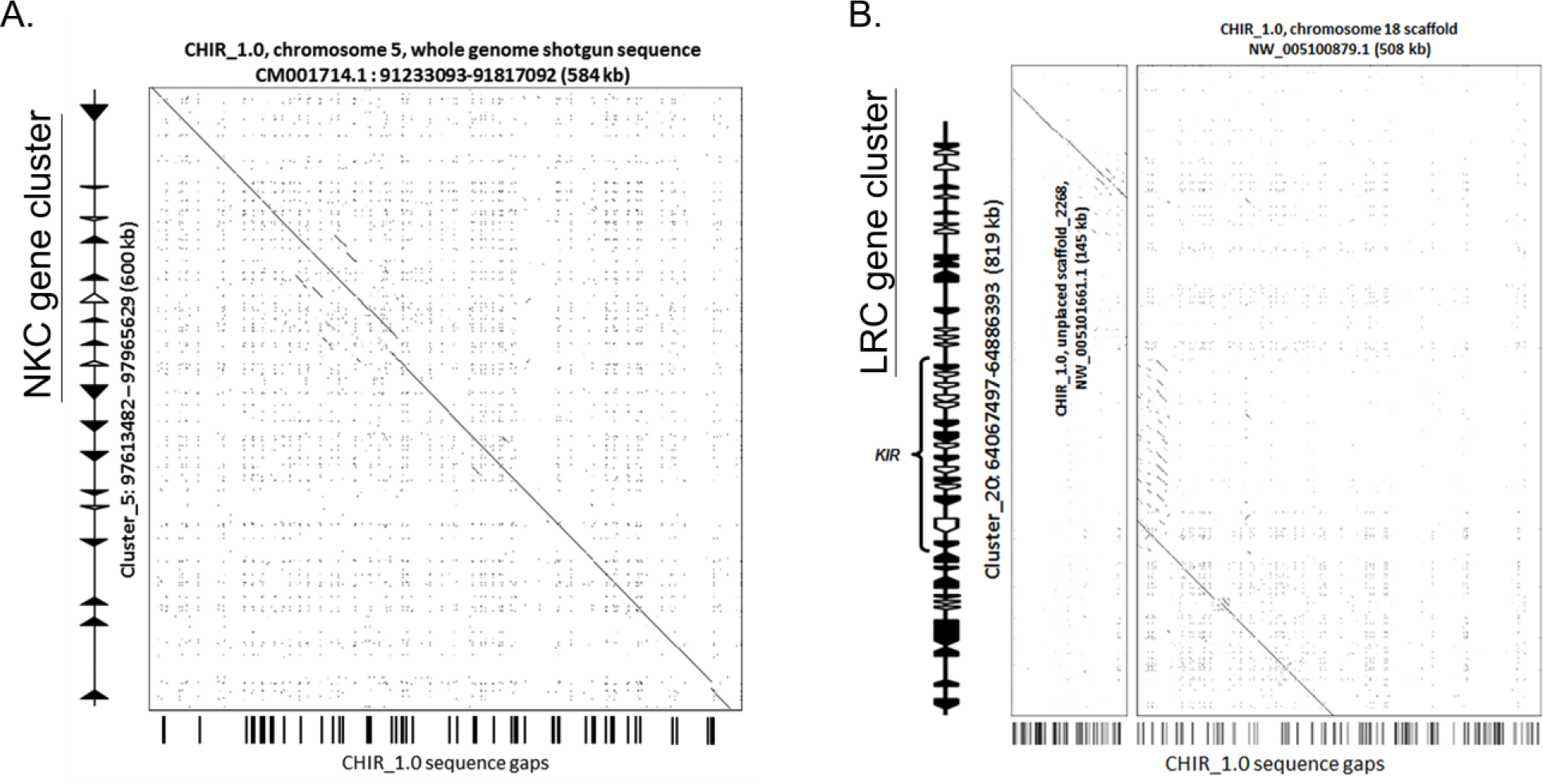
(A) A region of the Natural Killer Cell (NKC) gene cluster was highly fragmented in the CHIR_1.0 reference genome but is present on a single contig within ARS1. (B) Likewise, the Leukocyte Receptor Complex (LRC) locus was poorly represented in CHIR_1.0, and was missing ~600 kb of sequence. For highly repetitive and polymorphic gene families, our assembly approach provided the best resolution and highest continuity of gene sequence. Comparisons of our assembly to CHIR_2.0 show that this assembly is also missing this same intervening portion of sequence within the LRC.

### Structural elements and karyotype

Our assembly significantly improves on repeat resolution versus previous assembly approaches, including both short-read and Sanger sequencing projects^44, 45^. While not fully complete, we include large fractions of the allosomes and heterochromatic regions, which are typically absent from *de novo* assembly efforts. For example, we assembled >5 kbp of telomeric sequence on six autosomes, and additional, low complexity sequence on seven autosomes that was not present in CHIR_1.0 or CHIR_2.0. Using previously determined centromeric repeat sequence for goat^46^, we identified 15 of our chromosome scaffolds that included centromeric repeats greater than 2 kbp in length (Methods). Seven of these chromosomes (chromosomes 1, 6, 12, 13, 22, 26, and 29) had centromeric repeat sequence alignments that were larger than 8 kbp in length. Of our assembled chromosomes, we believe that we have assembled into both the centromere and telomeres on chromosomes 19 and 23, thereby capping the assembled chromosomes with constitutive heterochromatin. Two scaffolds (corresponding to chromosomes 13 and 28) have centromeric repeats 3 Mbp from the end, indicating the potential assembly of the elusive *p* arm of these acrocentric chromosomes (Methods). Additionally, closer examination of the optical maps revealed 34 maps containing large tandem and interspersed repetitive nickase motifs with a cumulative size of four megabases that did not align to the long-read contigs (Supplementary Table S6). Given that these repetitive maps also did not align to any prior *Capra hircus* assembly, it is possible that these represent portions of constitutive heterochromatin that could not be assembled using other technologies. Using the optical map, we identified 105 additional repetitive patterns greater than 12 kb that were represented in our final assembly and were distributed among all Hi-C chromosome scaffolds with the exceptions of chromosomes 9 and 10. Finer scale repeat identification using the RepeatMasker^47^ algorithm revealed that we were able to resolve more of the larger classes of repetitive elements (greater than 1 kb in length) in our assembly due to longer contig sizes (Fig. 4b). Specifically, we assembled 66% more nearly complete (> 75% sequence length) BovB LINE repeats than CHIR_2.0 and found that 43.6% of the CHIR_2.0 gaps that ARS1 successfully closes coincided with BovB repeats greater than 3.5 kbp in length (Supplementary Fig. S6 and Supplementary Table S7).

We also attempted to resolve the goat sex chromosomes using the final data generated by successive scaffolding efforts. Our final ARS1 assembly contained two scaffolds that mapped to two different—but contiguous—regions of the X chromosome; representing 85.9% of the expected chromosome size (based on a hypothesized X chromosome size of approximately 150 Mbp^39^). For the Y chromosome, self-hit alignment filtering and cross-species alignment to existing Y chromosome scaffolds in cattle identified 10 megabases of sequence that may have originated from the *Capra hircus* Y chromosome, approximately 50% of the estimated size^48^ (Supplementary Note 4 and Supplementary Table S8). Alignments of X-degenerate Y genes^49, 50^ and *Bos taurus* Y genes to these scaffolds confirmed their association with the Y chromosome, identifying 16% and 84% of our previously filtered contig list, respectively, with several contigs containing both sets of alignments. Both the heterochromatic nature of the Y chromosome and the ambiguous nature of the pseudoautosomal (the last portion of our X chromosome, and unplaced scaffolds 8, 12, 119 and 186) region’s placement on the X or Y chromosome precluded our ability to generate chromosome-scale scaffolds for this chromosome.

## Discussion

The advent of long-read sequencing has dramatically improved the average and N50 contig lengths of mammalian genome assemblies ^27, 31^, but complex genomic regions still interfere with the generation of complete, single contig chromosomes^31^. This is an even greater problem for polyploid genomes and species that have undergone whole genome duplications^51^. Therefore, reliable and affordable scaffolding technologies are vitally important for generating high-quality finished reference genome assemblies. In this study, we assessed the utility of both optical and chromatin interaction mapping, and found them to be complementary and most powerful in combination with long, high-quality PacBio contigs. As in past assembly projects, we opted for a stepwise combination of these methods that leveraged their unique benefits to generate a final assembly.

Optical mapping yielded higher resolution and the resulting scaffolds were easier to validate than the Hi-C scaffolds, due to fewer conflicts with the PacBio contigs. However, we found that optical mapping was insufficient to generate full chromosome-scale scaffolds, with one notable exception being the single scaffold spanning goat chromosome 20 (Fig. 2b). Currently, the primary limitation of the type of optical mapping used in this study appears to be double-strand breaks caused by close, opposing nickase sites, which subsequently breaks the map assembly due to a lack of spanning optical reads. Optical map scaffolding generated only three confirmed assembly errors (3 / 336, or 0.9% of scaffolds), two of which were difficult to detect without the use of the RH map. Scaffolding with Hi-C was able to accurately assign contigs to their respective chromosome groups, as supported by our RH map data, 99.8% of the time; however, there were several orientation errors detected after contig ordering. This problem can be lessened with larger input contigs or the selection a different restriction enzyme during the initial Hi-C library construction. Contigs and scaffolds with low orientation quality scores were frequently associated with orientation mistakes in the Hi-C scaffolds (r = 0.49; Pearson’s correlation) (Supplementary Table S2), suggesting that more frequent cutting may provide higher fidelity association results in the future. Regardless, recent advances in both methods achieved here the reconstruction of 29 vertebrate autosomes into single scaffolds with a minimal number of gaps (477) and without manual finishing.

Additionally, our assembly has improved the resolution of the goat sex chromosomes; however, complete assembly is complicated by the decreased coverage of these haploid chromosomes in the male individual combined with the complex distribution of heterochromatin and repetitive elements. Recent efforts to assemble the Y chromosomes of other species have relied on Y-associated BAC sequencing^52^ or chromosome sorting^53^ to increase the effective coverage. Here, Hi-C scaffolding was successful at clustering sex-chromosome contigs, but was unable to scaffold the Y chromosome or segregate X and Y chromosome contigs into singular distinctive clusters. Optical mapping also encountered difficulty in generating Y chromosome scaffolds, generating 16 scaffolds that contained 50.2% of the putative Y chromosome sequence in our assembly. Much of the sequence of the Y is constitutive heterochromatin^39^, which makes the generation of large optical maps and Hi-C fragments difficult.

Despite significant improvements over previous assembly approaches, limitations remain in our approach. ARS1 is a haplotype-mixed representation of a diploid animal. Haplotype phasing is possible using single-molecule^54^ and Hi-C^55^ technologies, so a future aim is to generate a phased reference assembly. Secondly, the majority of constitutive heterochromatin, including centromeres and telomeres, as well as large tandem repeats, such as the nucleolus organizer regions, remain unresolved even in the human reference genome, which has undergone years of manual finishing. These are likely to remain unresolved unless sequence read lengths continue to increase in size in order to completely span these repetitive regions. Still, we note that our assembly shows a marked improvement at resolving the full structure of large repetitive elements, such as BovB retro-transposons and centromeric repeats (Fig. 4b). This increased resolution will enable future, pan-ruminant analysis of these repeat classes which may lead to further insight into the evolution of ruminant chromosome structure.

The methods presented in this study have generated chromosome-scale scaffolds, thereby reducing the extensive cost of genome finishing. By using a tiered approach to scaffold highly continuous single-molecule contigs, we obviate the need for expensive cytometry or BAC-walking experiments for chromosome placement. We estimate a current project cost in the neighborhood of $100,000 to complete a similar genome assembly, using RSII sequencing and the same scaffolding platforms used here. This cost is on the order of three times greater than a short read assembly scaffolded in a similar fashion, but with a tremendous gain in continuity and quality. To achieve similar quality via manual finishing of a short-read assembly would be cost prohibitive. Moreover, advances in single molecule sequencing including an updated SMRT platform and alternative nanopore-based platforms, will continue to decrease this cost in the near future to the point where generation of high-quality, chromosome-scale reference genome assemblies will approach the cost of a short-read assembly. As shown by the completeness of our assembly, and the improvements in gene model contiguity, we expect these methods will enable the scaling of *de novo* genome assembly to large numbers of vertebrate species without requiring major sacrifices with respect to quality.

## Methods

### Reference individual selection

A DNA panel composed of 96 U.S. goats from 6 breeds (35 Boer, 11 Kiko, 12 LaMancha, 15 Myotonic, 3 San Clemente, and 20 Spanish) was assembled to identify the most homozygous individual as the best candidate for genome sequencing. Genotypes were generated using Illumina’s Caprine53K SNP beadchip processed through Genome Studio (Illumina, Inc. San Deigo, CA). The degrees of homozygosity of individuals were determined by raw counts of homozygous markers on the genotyping chip. Individuals were ranked by their counts of homozygous markers and the individual with the highest count was selected as the reference animal.

### Genome sequencing, assembly, and scaffolding

Libraries for SMRT sequencing were constructed as described previously^31^ using DNA derived from the blood of the reference animal. We generated 465 SMRTcells using P5-C3 (311 cells), P4-C2 (142 cells), and XL-C2 (12 cells) chemistry (Pacific Biosciences). A total of 194 Gbp (69X) of subread bases with a mean read length of 5,110 bp were generated.

The “Celera Assembler PacBio corrected Reads” (CA PBcR) pipeline^30^ was used for assembly. Celera Assembler v8.2 was run with sensitive parameters specified in Berlin et. al ^30^ which utilized the MinHash Alignment Process (MHAP) to overlap the PacBio-reads to themselves and PBDAGCON^28^ to generate consensus for the corrected sequences. This generated 7.4 million error corrected reads (~38 Gbp; 5.1 kb average length) for assembly, which in turn produced 3,074 contigs having an NG50 of 4.159 Mbp with a total length of 2.63 Gbp and 30,693 degenerate contigs <50 kbp in length with a total length of 288.361 Mbp. Initial polishing was performed with Quiver^28^ using the P5-C3 data only. The degenerate contigs (representing 9.90% of the 2.914 Gbp assembled length) were excluded from scaffolding by optical maps and Hi-C and incorporated into ARS1 as unplaced contigs.

Scaffolding of the contigs with optical mapping was performed using the Irys optical mapping technology (BioNano Genomics). DNA of sufficient quality was unavailable from the animal sequenced due to his accidental death, so we extracted DNA derived from a male child of the original animal. Purified DNA was embedded in a thin agarose layer was labeled and
counterstained following the IrysPrep Reagent Kit protocol (BioNano Genomics) as in Hastie et al.^21^. Samples were then loaded into IrysChips and run on the Irys imaging instrument (BioNano Genomics). The IrysView (BioNano Genomics) software package was used to produce singlemolecule maps and *de novo* assemble maps into a genome map. A 98X coverage (256 Gbp) optical map of the sample was produced in two instrument runs with labeled single molecules above 100kb in size.

Scaffolding was also performed using Hi-C based proximity guided assembly (PGA). Hi-C libraries were created from goat WBC as described^56^, in this case the sequenced animal was used as samples were taken prior to his death. Briefly, cells were fixed with formaldehyde, lysed, and the crosslinked DNA digested with HindIII. Sticky ends were biotinylated and proximity-ligated to form chimeric junctions, that were enriched for and then physically sheared to a size of 300-500 bp. Chimeric fragments representing the original crosslinked long-distance physical interactions were then processed into paired-end sequencing libraries and 115 million 100bp paired-end Illumina reads were produced. The paired-end reads were uniquely mapped onto the draft assembly contigs which were grouped into 31 chromosome clusters, and scaffolded using Lachesis software^15^ with tuned parameters discovered by Phase Genomics (Supplementary Note 1).

### Conflict resolution

The tiered approach to scaffolding that we used provides several opportunities for the resolution of misassemblies in the initial contigs, and contig orientation mistakes made by the various methods of scaffolding, respectively. In order to resolve such conflicts, we used a consensus approach that used evidence from six different sources of information: (a) a previously generated RH map^34^, (b) our long read-based contig sequence, (c) Irys optical maps, (d) Hi-C scaffolding orientation quality scores, and (e) Illumina HiSeq read alignments to the contigs (Fig. 1b). We found that 40 contigs did not align with the Irys optical map, and there were 102 Irys conflicts that needed resolution. A large proportion of the conflicts were identified as forks in the minimum tiling path of contigs superimposed on Irys maps, but we found that 70 of these conflicts were due to ambiguous contig alignments on two or more Irys maps. These ambiguous alignments were due to the presence of segmental duplications on multiple scaffolds, and were discarded. Of the original 102 conflicts, only 32 conflicts had characteristic drops in Illumina sequence read depth and RH map order conflicts that were indicative of mis-assembly. The intial Irys-PGA dataset had 220 orientation conflicts with our previously generated RH map. Approximately 41.8% of these orientation conflicts were resolved by PBJelly gap filling at alternative scaffold junctions (92 / 220) suggested by the RH map. Unlike the optical map-contig conflict resolution step, we were unable to identify a suitable consensus model that enabled us to predict and resolve all 220 orientation conflicts without generating a large number of false positive scaffold breaks. Since the *Capra hircus* X chromosome is acrocentric, our two X chromosome scaffolds do not represent distinct arms of the goat X chromosome and were likely split due to the requested number of clusters in the proximity-guided assembly algorithm. Still, our recommendation is to use the haploid chromosome count as input to Hi-C scaffolding to avoid false positive scaffold merging. We recommend the use of a suitable genetic or physical map resource, the use of larger input scaffolds into the PGA algorithm or the use of more frequent cutting restriction enzymes in the generation of Hi-C libraries in order to avoid or resolve these few remaining errors.

### Assembly polishing and contaminant identification

After scaffolding and conflict resolution we ran PBJelly from PBSuite v15.8.24^36^ with all raw PacBio sequences to close additional gaps. PBJelly closed 681 of 1,439 gaps of at least 3 bp in length. A final round of Quiver was run to correct sequence in filled gaps. It removed 846 contigs with no sequence support, leaving 663 gaps. Finally, as P5-C3 chemistry is higher error than either P4-C2 or P6-C4 (https://github.com/PacificBiosciences/GenomicConsensus/blob/master/doc/FAQ.rst), we generated 23X coverage of the initial goat sample using 250 bp insert Illumina HiSeq libraries for post-processing error correction. We aligned reads using BWA^57^ (version: 0.7.10-r789) and Samtools^58^ (version: 1.2). Using PILON^37^, we closed 1 gap and identified and corrected 653,246 homozygous insertions (885,794 bp), 87,818 deletions (127,024bp), and 34,438 (34,438bp) substitutions within the assembly that were not present in the Illumina data. This matches the expected error distribution of PacBio data, which has ~5-fold more insertions than deletions^59^. Closer investigation of this data revealed that the majority of insertion events (52.01%) were insertions within a homopolymer run, a known bias of the PacBio chemistry [https://github.com/PacificBiosciences/GenomicConsensus/blob/master/doc/FAQ.rst]. PILON also identified 1,082,330 bases with equal-probability heterozygous substitutions, indicating potential variant sites within the genome.

The final assembly was screened for viral and bacterial contamination using Kraken v0.10.5^60^ with a database including Viral, Archeal, Bacterial, Protozoa, Fungi, and Human. A total of 183 unplaced contigs and 1 scaffold were flagged as contaminant and removed. An additional two unplaced contigs were flagged as vector by NCBI and removed.

### Assembly annotation

We employed EVidence Modeler (EVM)^61^ to consolidate RNA-seq, cDNA, and protein alignments with *ab initio* gene predictions and the CHIR_1.0 annotation into a final gene set. RNA-seq data included 6 tissues (hippocampus, hypothalamus, pituitary, pineal, testis, and thyroid) extracted from the domesticated San Clemente goat reference animal and 13 tissues pulled from NCBI SRA (Supplementary Table S9). Reads were cleaned with Trimmomatic^62^ and aligned to the genome with Tophat2^63^. Alignments were then assembled independently with StringTie^64^, Cufflinks^65^ and *de novo* assembled with Trinity^66^. RNA-seq assemblies were then combined and further refined using PASA^61^. Protein and cDNA alignments using exonerate and tblastn with Ensembl datasets of *Ovis aries, Bos taurus, Equus caballus, Sus scrofa,* and *Homo sapiens* as well as NCBI annotation of *Capra hircus* and *Ab initio* predictions by Braker1^67^ were computed. The CHIR_1.0 annotation coordinates were translated into our coordinate system with the UCSC lifOver tool. All lines of evidence were then fed into EVM using intuitive weighting (RNAseq > cDNA/protein > *ab initio* gene predictions). Finally, EVM models were updated with PASA.

### Gap resolution and repeat analysis

Sequence gap locations were identified from the CHIR_1.0, CHIR_2.0 and ARS1 assembly using custom Java software (https://github.com/njdbickhart/GetMaskBedFasta). In order to identify identical gap regions on different assemblies, we used a simple alignment heuristic. In brief, we extracted 500 bp fragments upstream and downstream of each gap region in CHIR_1.0 or CHIR_2.0 and then aligned both fragments to the assembly of comparison (eg. ARS1). If both fragments aligned successfully within 10 kb, which was a length greater than 99.6% of all CHIR_1.0 and CHIR_2.0 gaps, on the same scaffold/chromosome and the intervening sequence did not contain ambiguous (N) bases, the gap was considered “closed.” If fragments aligned to two separate scaffolds/chromosomes, then the region was considered a “trans-scaffold” break. In cases where one or both fragments surrounding a gap did not align, or if there were two or more ambiguous bases between aligned fragments, the gap was considered “open.” Repeats were identified using the RepBase library release 2015-08-07 with RepeatMasker^47^ on the ARS1, CHIR_2.0, UMD3.1 (cattle)^44^ and Oarv3.1 (sheep; http://www.livestockgenomics.csiro.au/sheep/oar3.1.php) reference assemblies. The “quick” (-q) and “species” (e.g.,-species goat,-species sheep,-species cow) options were the only deviations from the default. Repeats were filtered by custom scripts if they were less than 75% of the expected repeat length or were below 60% identity of sequence. Gap comparison images between assemblies were created using Nucmer^68^.

### Centromeric and telomeric repeat analysis

To identify telomeric sequence we used the 6-mer vertebrate sequence (TTAGGG) and looked for all exact matches in the assembly. We also ran DUST^69^ with a window size of 64 and threshold of 20. Windows with at least 10 consecutive identical 6-mer matches (fwd or rev strand) intersecting with low-complexity regions of at least 1500 bp were flagged as potential telomeric sites and those with >5 kbp total length reported. To identify putative centromeric features in our assembly, we used centromeric repetitive sequence for goat from a previously published study^46^. Subsequent alignments of that sequence were used to flag collapsed centromeric sequence in our assembly, identifying three unplaced contigs that contained large portions of the repeat. The contigs were mapped to the assembly and regions at least 2 kbp in length reported as centromeric sites. In all but four cases the telomeric and centromeric sequences were within 100 kbp of the contig end (Supplementary Table S10). In the cluster corresponding to chromosome 1, the centromeric sequence was at position 40 Mbp, confirming a mis-assembly identified by the RH map. In chromosomes 12 and 13 (clusters 13 and 14, respectively) the centromere was <3 Mbp from the end, indicating potential assemblies of the short chromosome arms, though this has not yet been experimentally confirmed

## Acknowledgments

We thank Robert Lee for technical assistance. This project was supported by the US Agency for International Development Feed the Future program, Norman Borlaug Commemorative Research Initiative, Livestock Improvement Program. This work was also supported in part by Agriculture and Food Research Initiative (AFRI) competitive grant No. 2011-67015-30183 and 2015-67015-22970 from the USDA National Institute of Food and Agriculture (NIFA) Animal Genome Program. DMB, BR, SGS, and CPVT were supported by USDA CRIS project number: 8042-31000-104-00. CMK and TPLS were supported by USDA CRIS project number: 3040-31320-012-00. GCW was supported by USDA CRIS project number: 6402-31000-006-00D. JS was supported in part by NIH R01 grant number HG006283. MJD and IL were supported in part by NIH grant number P41 GM103533. MJD is a Senior Fellow in the Genetic Networks program at the Canadian Institute for Advanced Research and a Rita Allen Foundation Scholar. IL is supported by the UW Commercialization Gap Fund and Commercialization Fellows Program. FAPL was supported by MN Experiment Station Project MIN-16-103. JCS and JAH were funded by the United Kingdom Biotechnology and Biological Sciences Research Council Institute Strategic Program on Livestock Viral Diseases awarded to The Pirbright Institute. Mention of trade names or commercial products in this article is solely for the purpose of providing specific information and does not imply recommendation or endorsement by the US Department of Agriculture. SK and AMP were supported by the Intramural Research Program of the National Human Genome Research Institute, National Institutes of Health. This study utilized the computational resources of the Biowulf system at the National Institutes of Health, Bethesda, MD (http://biowulf.nih.gov

### Contributions

DMB, BR, SK, TSS, GEL, JS, AMP, CPVT and TPLS planned and coordinated the study and wrote the manuscript. TPLS and CMK performed the long-read sequencing and assisted with downstream analysis. SK and AMP performed the initial long-read assembly. ARH, SC, JL, and ETL performed the optical mapping and provided technical support related to the data. IL, STS, JB, MJD, and JS designed the Hi-C experiments, produced assembly scaffolds from the data and provided technical support. DMB and SK polished the final reference assembly. JAH and JCS provided manual annotation of the immune gene clusters. FAPDL provided manual annotation of the Y chromosome genes and contigs. BLS and Jiajie S extracted RNA-seq biopsies and ran RNA-seq experiments, respectively. BLS, JLH, YZ, Jiajie S, HJH, GCW, and AC performed downstream analysis of the data and assisted in the generation of additional files for the manuscript. All authors read and approved the final manuscript.

### Competing Financial Interests

TSS is a current employee of Recombinetics. IL and STS are employees of Phase Genomics. JB, JS and MJD have a vested financial interest in Phase Genomics. ARH, SC, JL, and ETL are employees of BioNano Genomics. All other authors declare no competing financial interests.

### Accession Codes

BioProject accession for all assembly datasets derived from the San Clemente goat reference animal: PRJNA290100. The ARS1 assembly: LWLT00000000. The Pacbio reads (SRA: SRP069238) the Illumina WGS reads (SRA: SRX1890394) and the Hi-C library reads (SRA: SRX1910977) were uploaded to public databases. RNA-seq reads were uploaded to the SRA (Accession: SUB1684327). Currently, there are no databases for storing optical mapping data, so this data is available at the following link: https://gembox.cbcb.umd.edu/goat/index.html.

